# Classification of Indian States and Union Territories based on their Soil Macronutrient and Organic Carbon Profiles

**DOI:** 10.1101/2020.02.10.930586

**Authors:** Prashant Kaushik

## Abstract

Soil fertility determines the successful development of a plant, and therefore it is important to achieve food security. Imbalanced and inadequate use of chemical fertilizers, irregular irrigation and harmful cultural practices deplete the soil profile nutrient profile, which is critical for the successful crop production. This study presents the results of the classification of the states across India based on their soil macronutrient profile. The entanglement coefficient of Unweighted Pair Group Method with Arithmetic Mean (UPGMA) and the neighbour-joining method was 0.81. Absolute correlation values were determined among the different classes of the soil nitrogen content and the corresponding classes of the soil phosphorus content. The K-mean clustering method divided the states and union territories into the three clusters. Overall, this works represents the grouping of Indian soils based on their soil macronutrient and organic carbon content.

## Introduction

Soil fertility is critical for successful crop production. As the population is continuously rising, there is pressure to produce more food. Continuous cropping for enhanced yield removes substantial amounts of nutrients from the soil. Soil fertility is usually determined based on the presence or absence of nutrients, i.e. macro and micronutrients^1,2^. Out of 17 critical plant nutrients Nitrogen (N), Phosphorus (P), potassium (K), calcium (Ca), magnesium (Mg), and sulphur (S) are macronutrients. The sustainable productivity of soil depends upon its potential to provide critical nutrients to plants. Therefore, evaluation of the fertility status of soils is an essential aspect of sustainable agriculture^3^.

In this direction soil health card scheme launched by the Government of India in 2015 has provided valuable information regarding the soil profile of Indian states across India. In this scheme, the farmers are provided with the soil health cards to the farmers to provide them with the crop-wise suggestions to enhance productivity employing judicious use of fertilizers for achieving yield targets. The soil profile was provided based on the soil testing for the macronutrients in the various labs settled across India. The outcome and suggestion are displayed inside the cards known as the soil health card that is also available online^4,5^.

Soil testing assesses the fertility status and delivers information and facts with regards to nutrient availability in soils. This information is the basis for the fertilizer suggestions for maximizing crop yields. For right soil management, the farmer should know what amendments are essential to optimize the productivity of soil for the specific crops^6,7^. Here the characterization of the soils across Indian states and union territories based on their macronutrient profiles data available from the online database known as soil health card scheme of the government of India.

## Material and Methods

### Data and Classification

The data was taken from the soil health card scheme from the online repository https://soilhealth.dac.gov.in/.

Briefly, the data was generated as part of the Indian government initiative to started in 2015. The nitrogen, phosphorus, potassium and organic carbon content were determined based on the procedures defined elsewhere^8^. Further, the five-level classification of macronutrients presents in the Indian soils was undertaken similar to the one based on the method defined similarly to one defined elsewhere^9^.

**Table 1.** The five-tier classification of the N, P, K and organic carbon present in the Indian soils.

| **Macronutrient** | **Code** |
| --- | --- |
| Nitrogen Very Low | NVL |
| Nitrogen Low | NL |
| Nitrogen Medium | NM |
| Nitrogen High | NH |
| Nitrogen Very High | NVH |
| Phosphorus Very Low | PVL |
| Phosphorus Low | PL |
| Phosphorus Medium | PM |
| Phosphorus High | PH |
| Phosphorus Very High | PVH |
| Potassium Very Low | KVL |
| Potassium Low | KL |
| Potassium Medium | KM |
| Potassium High | NH |
| Potassium Very High | KVH |
| Organic Carbon Very Low | CVL |
| Organic Carbon Low | CL |
| Organic Carbon Medium | CM |
| Organic Carbon High | CH |
| Organic Carbon Very High | CVH |

### Study Area

The list of Indian states and union territories used in the present study is presented in the Table 2.

**Table 2.** List of Indian states and union territories used in the study.

| **State/Union Territory** | **Code** |
| --- | --- |
| Andaman and Nicobar Islands | AN |
| Andhra Pradesh | AP |
| Arunachal Pradesh | AR |
| Assam | AS |
| Bihar | BR |
| Chhattisgarh | CG |
| Dadra and Nagar Haveli and Daman and Diu (union territory) | DN |
| National Capital Territory of Delhi (union territory) | DL |
| Goa | GA |
| Gujarat | GJ |
| Haryana | HR |
| TerrHimachal Pradesh | HP |
| Jammu and Kashmir (union territory) | JK |
| Jharkhand | JH |
| Karnataka | KA |
| Kerala | KL |
| Madhya Pradesh | MP |
| Maharashtra | MH |
| Manipur | MN |
| Meghalaya | ML |
| Mizoram | MZ |
| Nagaland | NL |
| Odisha | OR |
| Puducherry (union territory) | PY |
| Punjab | PB |
| Rajasthan | RJ |
| Sikkim | SK |
| Tamil Nadu | TN |
| Telangana | TG |
| Tripura | TR |
| Uttar Pradesh | UK |
| Uttarakhand | UP |
| West Bengal | WB |

## Data Analysis

Data analysis was performed in the R environment using the N, P, K and organic carbon profile of different Indian states. In this direction, the data was subjected to Unweighted Pair Group Method with Arithmetic Mean (UPGMA) hierarchical clustering and ward’s clustering (R Core Team 2015). The “average” method was used with the function hclust. Further, the dendrograms were compared with the tanglegram algorithm using the R package dendextend^10^. Whereas, the correlation coefficients among the various classes of all of the four macronutrients were plotted using the packages corrplot ^11^. Finally, K-mean clustering was also performed in the R environment.

## Results and Discussion

Based on the UPGMA clustering, the states were branched into two main clusters and many sub-clusters (Figure 1). AP (Andhra Pradesh) and WB (West Bengal) were clustered together apart from the rest of the states and the union territories (Figure 1). Similarly, the union territory DN and state TR coexisted together (Figure 1).

**Figure 1.**
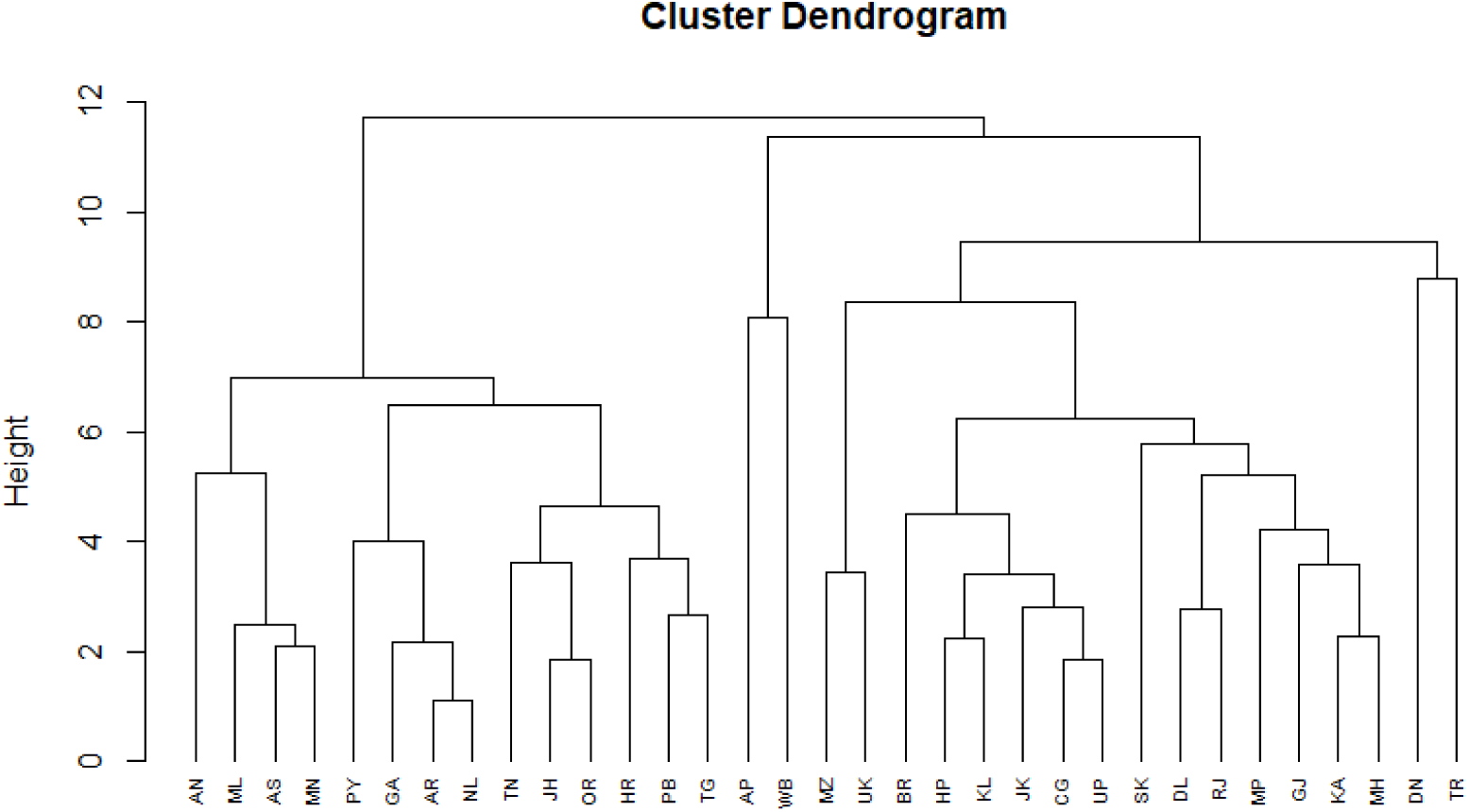
Unweighted Pair Group Method with Arithmetic Mean (UPGMA) clustering of Indian states based on their macronutrient profile.

The tangelgram is represented in Figure 2. The entanglement coefficient of Unweighted Pair Group Method with Arithmetic Mean (UPGMA) and the neighbour-joining method was 0.81 (Figure 2).

**Figure 2.**
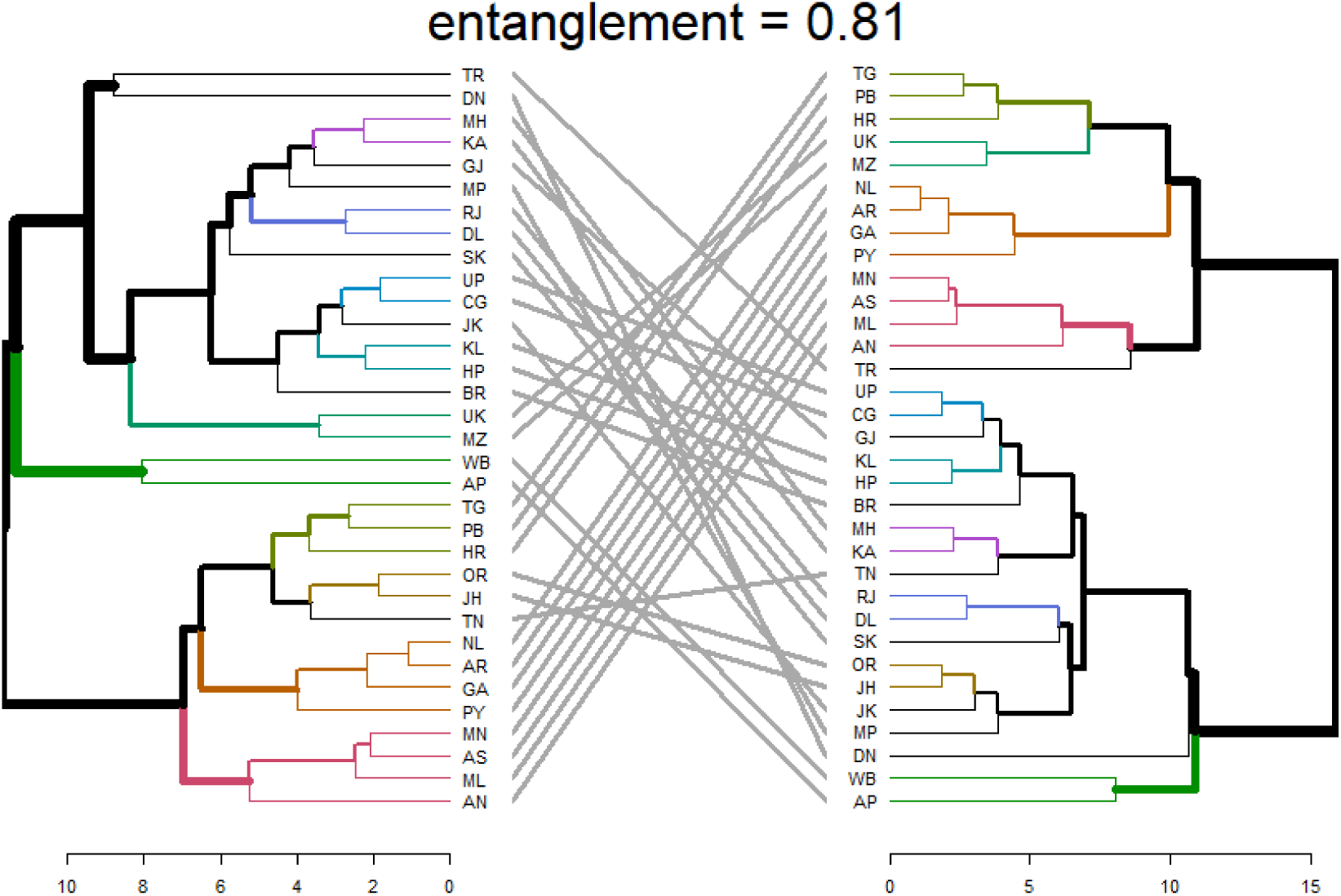
Tangelgram showing the relationship between the two different methods used for the clustering of Indian states UPGMA (Left) and neighbour-joining method (right).

Pearson’s correlations among the different categories of the N, P, K and organic carbon are represented in Figure 3. Five positive and absolute correlations were determined among the various classes of the soil nitrogen content and the corresponding classes of the soil phosphorus content (Figure 3).

**Figure 3.**
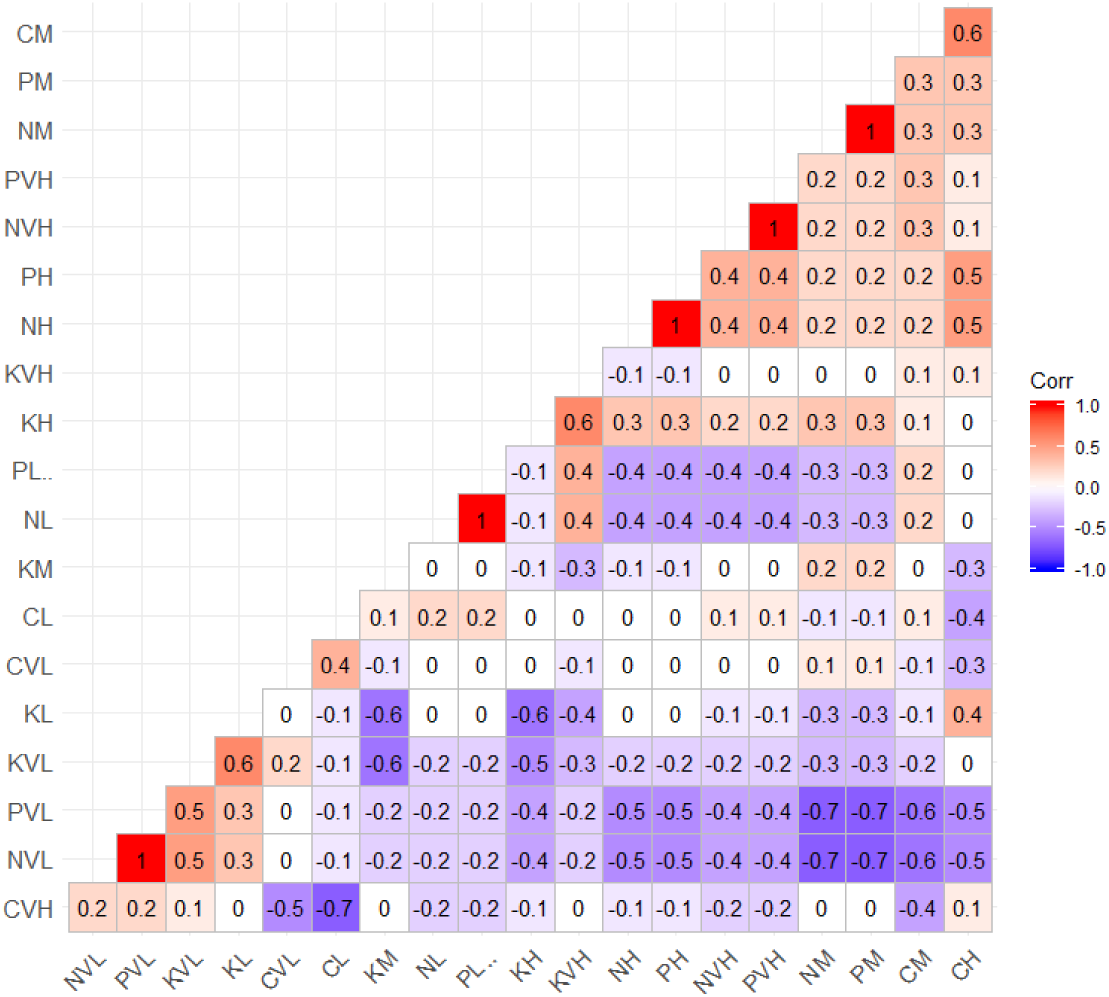
Pearson’s correlations among the six classes of the four macronutrients used for the classification of Indian states in this study.

There were three clusters based on the K-mean clustering results (Figure 4). The based on the soil micronutrient profiles states were unevenly divided irrespective of their geography (Figure 4). Interestingly, TN (Tamil Nadu) and TR (Tripura) were represented in the overlap zone of the clusters (Figure 4).

**Figure 4.**
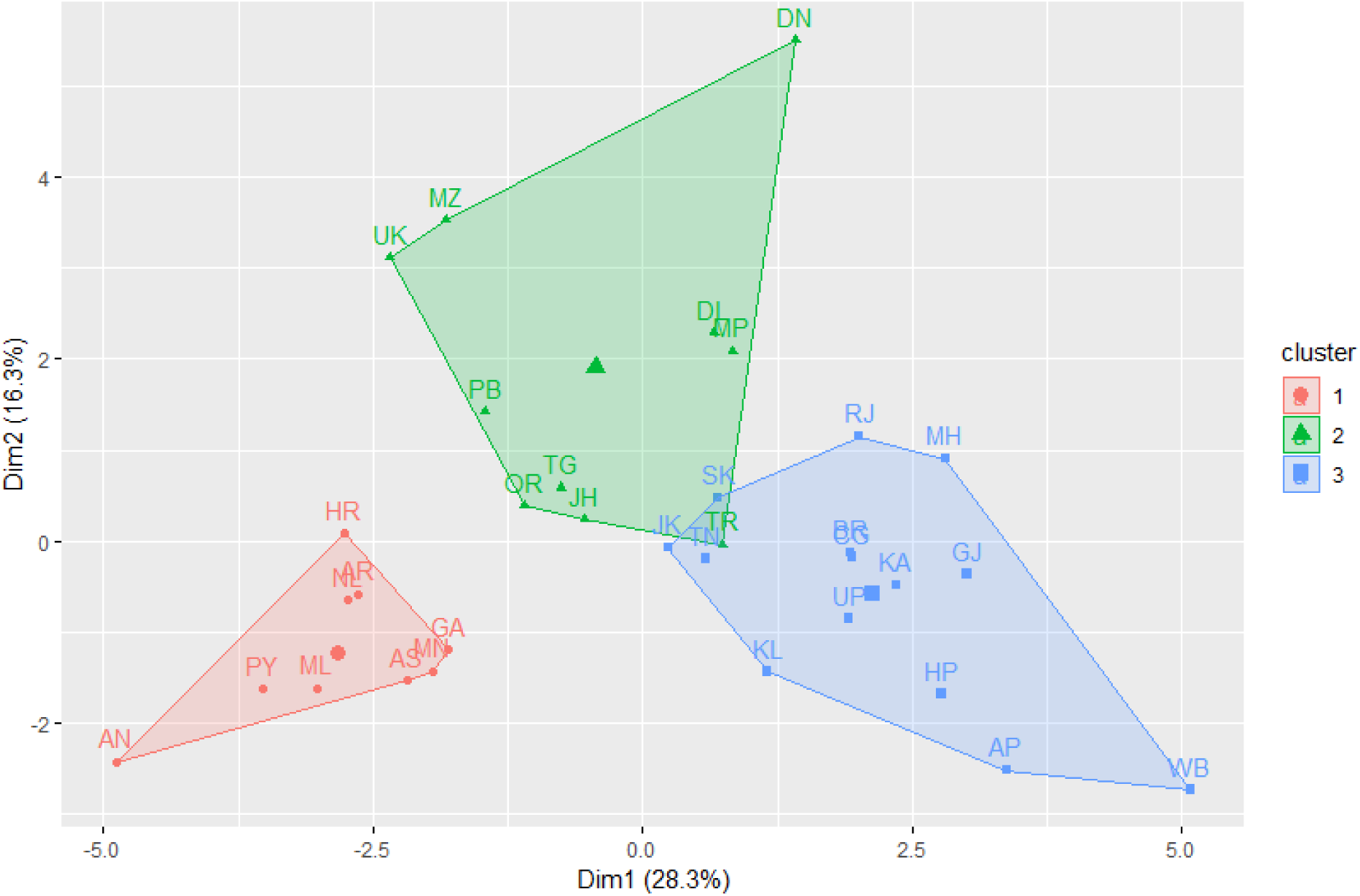
K-mean clustering of Indian states based on their macronutrient profile.

Recently, the amount of information regarding the soil classification based on their macronutrient and organic carbon profile has increased significantly^12^. Moreover, new technologies are also speeding up the same process with a more significant idea of environmental procedures. The classification algorithms are valuable to seek out subsets or styles of comparable samples determined by their data^13,14^. Even though many algorithms are circumscribed by several principles but the decision of the acceptable algorithm relies upon on conditions^15^. The method of K-mean clustering is extensively used in several research areas. Clustering method works based on the unsupervised learning approach; this means it works without any prior information regarding the data structure. The purpose of K-means clustering divides the samples with the initial values randomly selected from the data with updating of a grouping of data until a defined number of iterations are reached. With a first step of deciding the number clusters and later the initial centroids are randomly determined^16–18^.

